# Identifying Plasmids in Bacterial Genome Assemblies

**DOI:** 10.1101/332049

**Authors:** Barry G. Hall

**Affiliations:** Bellingham Research Institute Portland, OR 97212

**Keywords:** microbiology, pathogenesis, bioinformatics, plasmids, population biology

## Abstract

Despite their importance to bacterial pathogenesis, plasmids are rarely identified in incomplete genome sequences. The free FindPlasmids (FP) package for Windows, Mac OS X, and Linux facilitates identification of plasmids in incomplete genome sequences. FP found plasmids in 98.8% of complete genomes in which they were present, correctly identifying plasmids ranging in number from 1 to 10 plasmids. In a sample of 50 *E. coli* genome assemblies it has identified from zero to three plasmids ranging in size from 1,549 to 133,843 bp and present in 42% of the assemblies examined.

## Background

Plasmids play key roles in the ecology of bacteria, and are of particular interest to the medical community because they carry genes for antibiotic resistance, toxins, and virulence (Bennett 2008; Jackson et al. 2011; Gyles and Boerlin 2014). Over 12,000 plasmids have been identified, sequenced, and their genes described. The advent of Next Generation Sequencing has led to the sequencing of bacterial pathogens as an almost routine part of clinical studies, but in the vast majority of cases sequencing project are carried only to the genome assembly stage, making it difficult to determine what plasmids - and their antibiotic resistance and virulence determinants - are present in the sequenced genomes.

### Among 10,467 *Escherichia coli* genomes at the NCBI

(https://www.ncbi.nlm.nih.gov/genome/browse/#!/prokaryotes/Escherichia_coli) on May 20, 2018 there are 586 complete genomes. Of those 411, or 70.1%, include at least one plasmid. Among the 9881 incomplete genomes (scaffolds plus assemblies of contigs) only 53, or 0.54% are identified as containing a plasmid. Table 1 shows the frequencies of plasmids listed in complete and incomplete genomes of three additional species.

**Table 1.** Frequencies of plasmids being listed in complete and incomplete genomes

|  | Complete genomes |  | Incomplete Genomes |  |
| --- | --- | --- | --- | --- |
| Organism | Number | Percent with plasmids | Number | Percent indicated as containing plasmids |
| *Escherichia coli* | 586 | 70.1% | 9881 | 0.54% |
| *Pseudomonas aeruginosa* | 151 | 10.6% | 2615 | 0.076% |
| *Salmonella enterica* | 568 | 36.8% | 8115 | 0.099% |
| *Staphylococcus aureus* | 248 | 52.4% | 8046 | 0.44% |

While it is clearly rare to identify plasmid replicons in incomplete genomes, it is important to do so, especially in genomes of bacterial pathogens.

## FindPlasmids

Identification of plasmids in bacterial genome assemblies is facilitated by the FindPlasmids package which is freely available for Mac OS X, Linux, and Windows at https://sourceforge.net/projects/findplasmids/files/. FindPlasmids is based upon searching a local Blast+ database of plasmid sequences using a genome assembly as a query file. Detailed instructions for identifying a set of plasmids of interest, downloading those sequences, and making a local Blast+ database are included in the FindPlasmids package, as are command-line executable programs that facilitate making and searching the database. It typically takes about an hour to download the plasmid sequences and make a local Blast+ database.

The FindPlasmids program itself is a command-line executable that parses the results of the Blast+ search and identifies plasmids on the basis of the presence in the assembly of one or more contigs that match a plasmid sequence and whose total length constitutes at least 90% (user selectable) of the plasmid length. Plasmids are identified by GenBank accession number, making it easy to obtain the plasmid properties (coding sequences, etc.) from the corresponding GenBank file. It typically takes about a minute to identify the plasmids in a genome assembly.

To illustrate the use of FindPlasmids I made a Blast+ database of 5213 plasmids from members of the *Gamma Proteobacteria* (https://www.ncbi.nlm.nih.gov/genome/browse/#!/prokaryotes/plasmids,filtered_for_Gamma_Proteobacteria).

The FindPlasmids search returns a list of the plasmids that are hit by (match) any of the contigs in the sequence file. Plasmids are identified as likely being present in the genome if the sum of the hit lengths is at least 90% of the plasmid length. The mean fraction identical of those matches is reported, as are the identities of the contigs that hit the plasmid (Figure 1).

**Figure 1.**
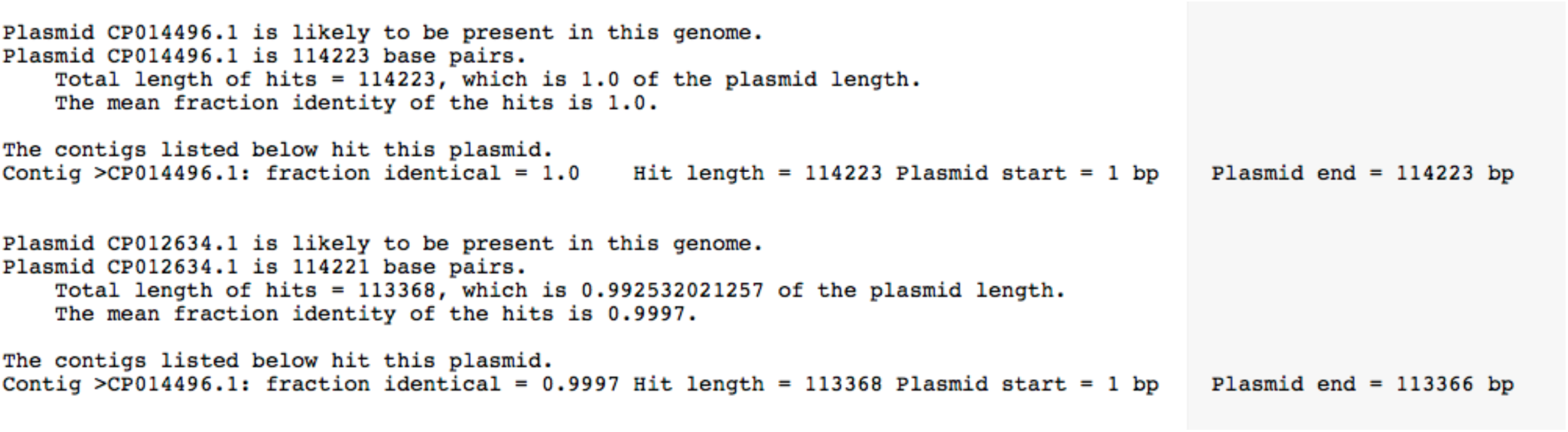
Result of searching the Gamma proteobacteria plasmids Blast+ database with the complete genome of *E. coli* strain SaT040.

First, it is important to understand that FindPlasmids cannot find a plasmid in a genome sequence unless that plasmid is included in the Blast+ database. Second, the set of Gamma Proteobacteria includes many closely related plasmids, and even several identical plasmids that have been given different accession numbers. As a result, the same contig or set of contigs may match several plasmids. When a contig matches several plasmids the plasmid that is matched over the greatest length of the plasmid and that has the highest sequence identity is the plasmid that is most likely to be present. In Fig. 1 two plasmids are shown as likely to be present: CP014496.1, matched by contig CP014496.1, and CP012634.1, also matched by contig CP014496.1. In this complete genome sequence contig CP014496.1 is just the plasmid sequence itself. Plasmid CP014496.1 is matched by the contig over 1.0 of the plasmid length, and with 1.0 sequence identity. Plasmid CP012634.1 is matched by the contig over only 0.9925 of its length and with only 0.9997 sequence identity. One would therefore conclude that plasmid CP014496.1 is present.

## Effectiveness of FindPlasmids

The only genome sequences in which we can be sure which plasmids are present are complete (closed) genomes. To assess the effectiveness of identifying the presence of plasmids in genome sequence files I searched the Gamma Proteobacteria plasmids Blast+ database with each of the 586 complete (closed) genome sequences. In closed genomes the contigs other than the chromosome are plasmids, thus correct matches will identify those plasmids as being present and matching the contig over its full length and with perfect sequence identity.

The number of plasmids per genome ranged from zero to 10, with a median of 1 and a mean of 1.76 plasmids per genome. 175 genomes had zero plasmids and 411 genomes had one or more plasmids.

There are several kinds of errors that might occur: (1) failure to find plasmids that are present (because those plasmids are not in the database), (2) identifying plasmids as being present when the contigs actually match part of the chromosome (3) identifying plasmids as being present that are not actually present.

FindPlasmids failed to find one or plasmids that were actually present in 5 genomes; i.e. in 0.85% of the complete genomes or 1.2% of the genomes that actually included plasmids. Zero plasmids that were identified were actually part of a chromosome. In 22 cases there were perfect matches to plasmids that were not present. In each case this was the result of the database including the same plasmid multiple times under different accession numbers. When searching incomplete genomes, where perfect matches are not expected, these redundant database entries would result in ambiguities; i.e. the same contig or set of contigs would match two plasmids over identical lengths and with the same sequences identity. Because those plasmids are the same it would not matter which was described as being present in that incomplete genome. FindPlasmids correctly identified the plasmids in 98.15% of genomes in which plasmids were present. It is reasonable to conclude that FindPlasmids is an effective and reasonably accurate tool for identifying plasmids in genome sequence files.

## Screening genome assemblies and scaffolds

50 *E. coli* genome assemblies that were not identified as containing plasmids were used to search the Gamma Proteobacteria Blast+ database. Of those 50 *E. coli* genomes 21, or 42%, carried close relatives of known plasmids (Table 2). Plasmid sizes ranged from 1,549 bp to 133,843 bp and were contained on from 1 to 122 contigs

**Table 2.**
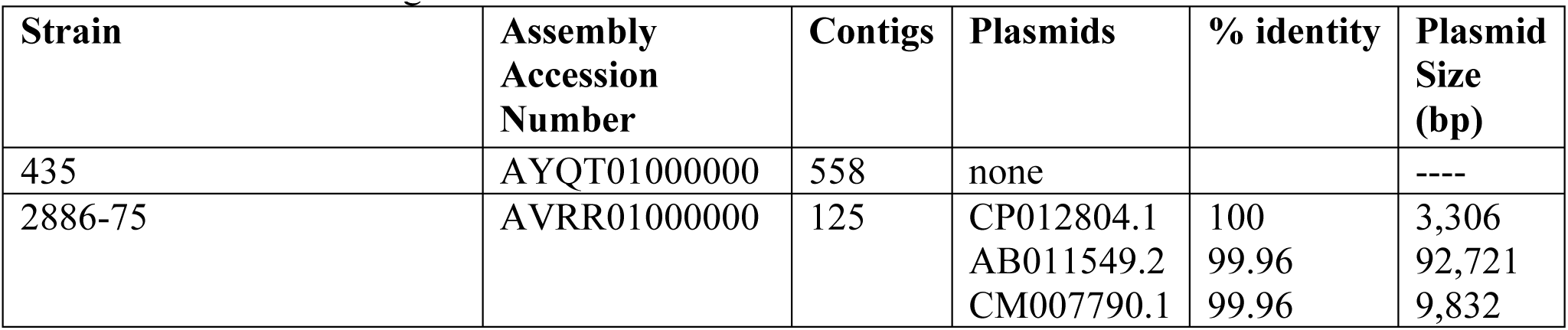
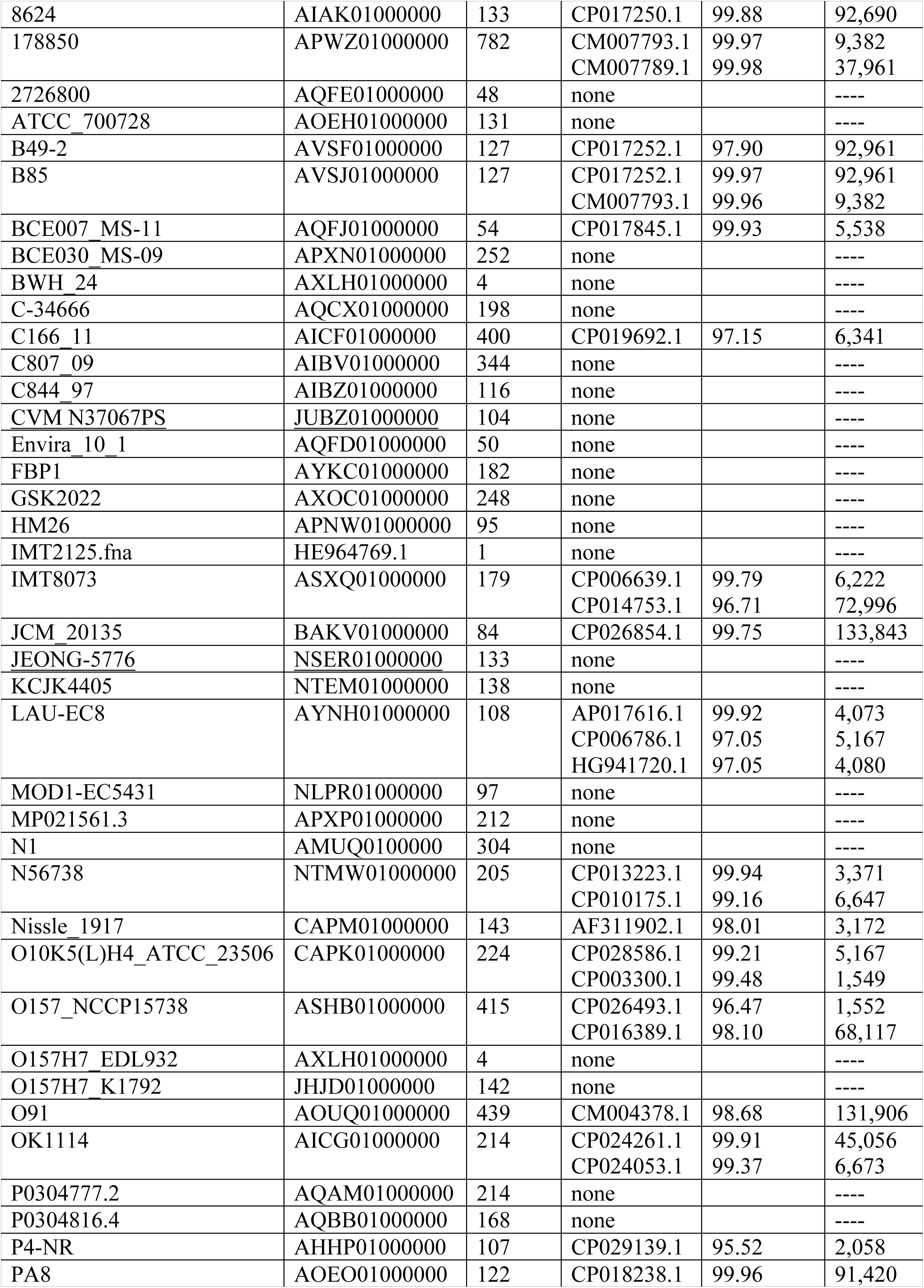
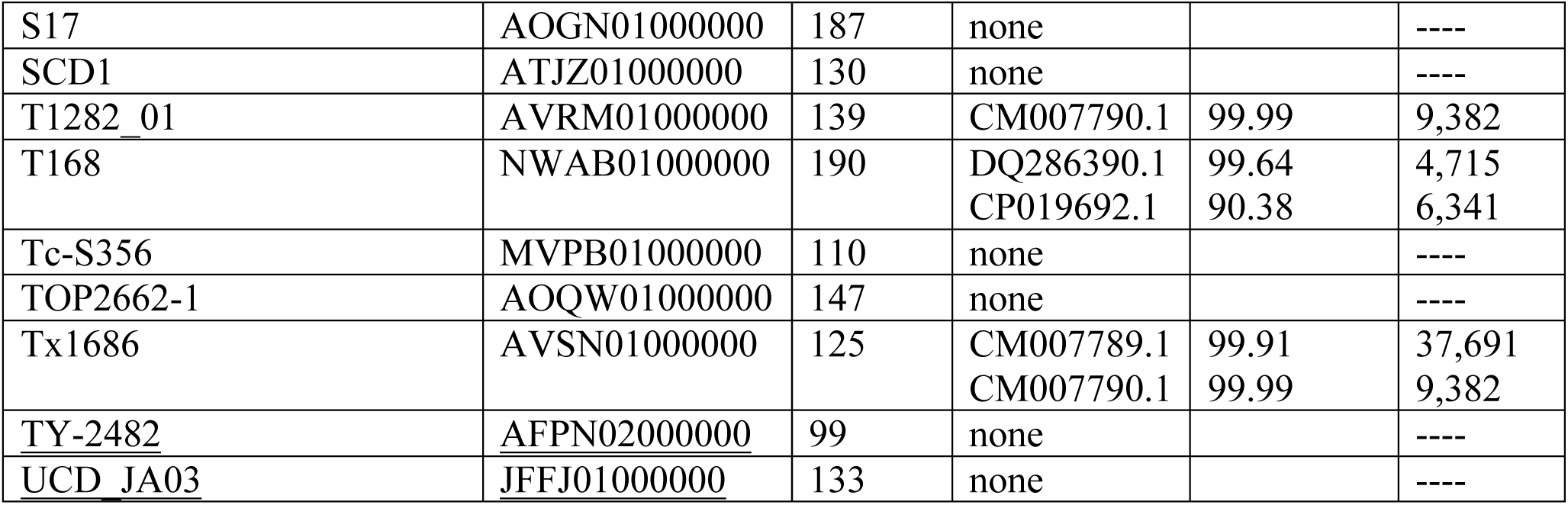
*Escherichia coli* genome assemblies

Table 3 shows that 40.3% of *E. coli* genome assemblies included one or more plasmids. That is a considerably smaller fraction than the 70.1% of complete genomes that contain plasmids but is about the same as in the 50 genome assemblies in Table 2. This suggests that complete genome sequences represent a biased sample of the *E. coli* population. That bias is probably based on the tendency to complete medically important genome sequences, many of which will carry virulence or antibiotic resistance determinants on plasmids. The same is true for *Pseudomonas aeruginosa* genome assemblies, where a bias toward plasmid-bearing strains is also evident in complete genomes. In contrast, for both *Salmonella enterica* and *Staphylococcus aureus*, the frequencies of plasmids in complete genomes and genome assemblies are about the same

**Table 3.**
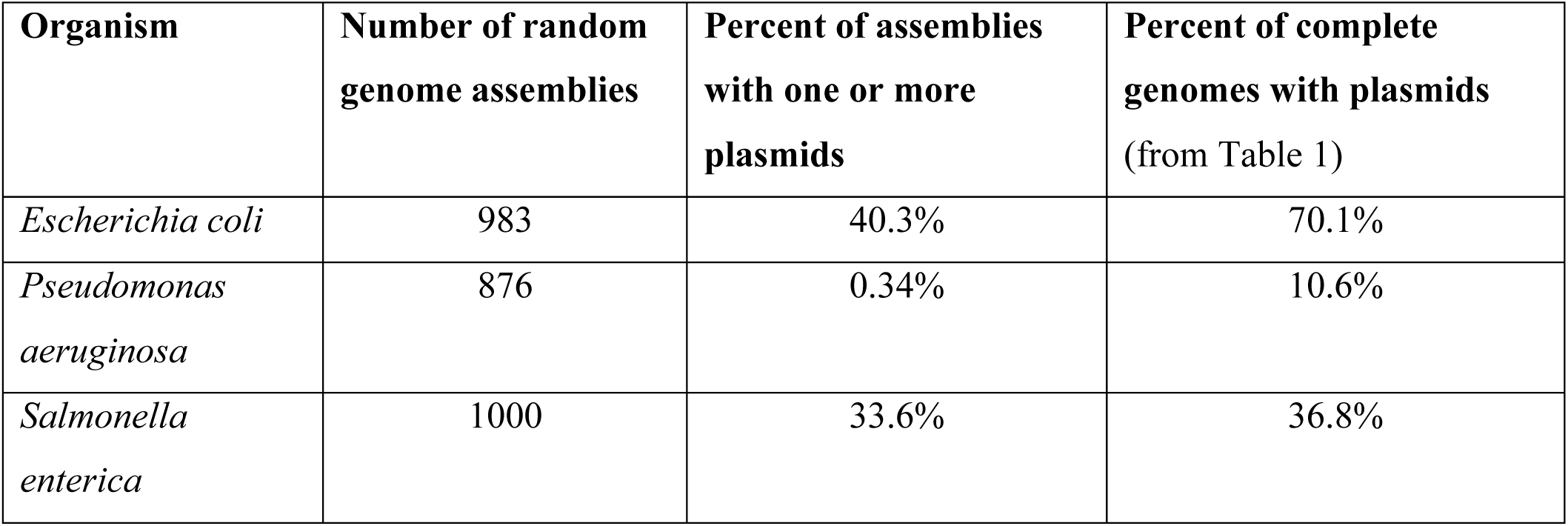
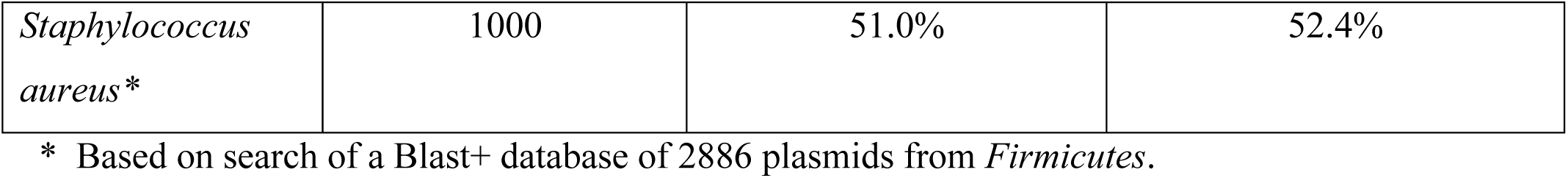
Plasmids in random samples of genome assemblies

| Organism | Number of random genome assemblies | Percent of assemblies with one or more plasmids | Percent of complete genomes with plasmids (from Table 1) |
| --- | --- | --- | --- |
| *Escherichia coli* | 983 | 40.3% | 70.1% |
| *Pseudomonas aeruginosa* | 876 | 0.34% | 10.6% |
| *Salmonella enterica* | 1000 | 33.6% | 36.8% |
| *Staphylococcus aureus* * | 1000 | 51.0% | 52.4% |

\* Based on search of a Blast+ database of 2886 plasmids from *Firmicutes*.

The FindPlasmids program appears to be an effective tool to detect the presence of plasmids in genome assemblies. It finds both small plasmids that constitute a single contig, and large plasmids that are distributed over many contigs. It detects the presence of plasmids in 98.8% of genomes where plasmids were present. That detection efficiency is strictly a function of the completeness of the plasmid Blast+ database. It is therefore important that the database err or the side of inclusion in order to detect broad host-range plasmids. Thus, even when looking for plasmids just in *E. coli* genomes it is better base the plasmid database on plasmids from the Gamma Proteobacteria than it is to base the database only on plasmids from *E. coli*.

Typically the identified plasmid is not identical to the contig(s) in the genome assembly, but is sufficiently similar to it to permit determining the presence of toxin genes, etc. (Table 2). If the plasmid is on a single contig it will be possible to submit the plasmid sequence to GenBank, obtain an accession number, and include that when the assembly is submitted to GenBank. Even when a plasmid is on several contigs, it should be possible to submit those contigs as an assembly of the plasmid sequence.

It is anticipated that routinely identifying plasmids in genome assemblies will facilitate studies of plasmid transmission and plasmid population genetics in collections of clinical isolates and will deepen our understanding of the roles of plasmids in bacterial pathogenesis.

## Acknowledgements

I am grateful to Miriam Barlow for pointing out the problem of identifying plasmids in genome assemblies.

